# Pleiotropy-guided transcriptome imputation from normal and tumor tissues identifies new candidate susceptibility genes for breast and ovarian cancer

**DOI:** 10.1101/2020.04.23.043653

**Authors:** Siddhartha P. Kar, Daniel P. C. Considine, Jonathan P. Tyrer, Jasmine T. Plummer, Stephanie Chen, Felipe S. Dezem, Alvaro N. Barbeira, Padma S. Rajagopal, Will Rosenow, Fernando M. Antón, Clara Bodelon, Jenny Chang-Claude, Georgia Chenevix-Trench, Anna deFazio, Thilo Dörk, Arif B. Ekici, Ailith Ewing, George Fountzilas, Ellen L. Goode, Mikael Hartman, Florian Heitz, Peter Hillemanns, Estrid Høgdall, Claus K. Høgdall, Tomasz Huzarski, Allan Jensen, Beth Y. Karlan, Elza Khusnutdinova, Lambertus A. Kiemeney, Susanne K. Kjaer, Rüdiger Klapdor, Martin Köbel, Jingmei Li, Clemens Liebrich, Taymaa May, Håkan Olsson, Jennifer B. Permuth, Paolo Peterlongo, Paolo Radice, Susan J. Ramus, Marjorie J. Riggan, Harvey A. Risch, Emmanouil Saloustros, Jacques Simard, Lukasz M. Szafron, Cheryl L. Thompson, Robert A. Vierkant, Stacey J. Winham, Wei Zheng, Jennifer A. Doherty, Andrew Berchuck, Kate L. Lawrenson, Hae K. Im, Ani W. Manichaikul, Paul D. P. Pharoah, Simon A. Gayther, Joellen M. Schildkraut

**Affiliations:** Medical Research Council Integrative Epidemiology Unit, University of Bristol, Bristol, UK; Population Health Sciences, Bristol Medical School, University of Bristol, Bristol, UK; Centre for Cancer Genetic Epidemiology, Department of Public Health and Primary Care, University of Cambridge, Cambridge, UK; Center for Bioinformatics and Functional Genomics, Department of Biomedical Science, Cedars-Sinai Medical Center, Los Angeles, CA; Section of Genetic Medicine, Department of Medicine, University of Chicago, Chicago, IL; Section of Hematology/Oncology, Department of Medicine, University of Chicago, Chicago, IL; Center for Public Health Genomics, University of Virginia, Charlottesville, VA; Hospital Clínico San Carlos, Madrid, Spain; Divison of Cancer Epidemiology and Genetics, National Cancer Institute, Bethesda, MD; Division of Cancer Epidemiology, German Cancer Research Center (DKFZ), Heidelberg, Germany; Cancer Epidemiology Group, University Cancer Center Hamburg (UCCH), University Medical Center Hamburg-Eppendorf, Hamburg, Germany; Department of Genetics and Computational Biology, QIMR Berghofer Medical Research Institute, Brisbane, Australia; Centre for Cancer Research, The Westmead Institute for Medical Research, The University of Sydney, Sydney, Australia; Department of Gynaecological Oncology, Westmead Hospital, Sydney, Australia; Gynaecology Research Unit, Hannover Medical School, Hannover, Germany; Institute of Human Genetics, University Hospital Erlangen, Erlangen, Germany; Friedrich-Alexander University Erlangen-Nuremberg, Comprehensive Cancer Center Erlangen, Erlangen, Germany; MRC Human Genetics Unit, MRC Institute of Genetics and Molecular Medicine, The University of Edinburgh, Western General Hospital, Edinburgh, UK; Laboratory of Molecular Oncology, Hellenic Foundation for Cancer Research, Aristotle University of Thessaloniki School of Medicine, Thessaloniki, Greece; Department of Health Science Research, Division of Epidemiology, Mayo Clinic, Rochester, MN; Department of Surgery, Yong Loo Lin School of Medicine, National University of Singapore and National University Health System, Singapore; Saw Swee Hock School of Public Health, National University of Singapore and National University Health System, Singapore; Department of Gynecology and Gynecologic Oncology, Kliniken Essen-Mitte/ Evang., Essen, Germany; Department of Gynecology, Center for Oncologic Surgery, Charité Campus Virchow-Klinikum, Berlin, Germany; Department of Gynecology and Obstetrics, Hannover Medical School, Hannover, Germany; Department of Virus, Lifestyle and Genes, Danish Cancer Society Research Center, Copenhagen, Denmark; Molecular Unit, Department of Pathology, Herlev Hospital, University of Copenhagen, Copenhagen, Denmark; The Juliane Marie Centre, Department of Gynecology, Rigshospitalet, University of Copenhagen, Copenhagen, Denmark; Department of Genetics and Pathology, International Hereditary Cancer Center, Pomeranian Medical University, Szczecin, Poland; Department of Genetics and Pathology, University of Zielona Góra, Zielona Góra, Poland; David Geffen School of Medicine, Department of Obstetrics and Gynecology, University of California at Los Angeles, Los Angeles, CA; Institute of Biochemistry and Genetics, Ufa Federal Research Centre of the Russian Academy of Sciences, Ufa, Russia; Department of Genetics and Fundamental Medicine, Bashkir State University, Ufa, Russia; Radboud Institute for Health Sciences, Radboud University Medical Center, Nijmegen, The Netherlands; Department of Gynaecology, Rigshospitalet, University of Copenhagen, Copenhagen, Denmark; Department of Pathology and Laboratory Medicine, University of Calgary, Foothills Medical Center, Calgary, Canada; Genome Institute of Singapore, Human Genetics, Singapore; Department of Obstetrics and Gynecology, Klinikum Wolfsburg, Wolfsburg, Germany; Division of Gynecologic Oncology, University Health Network, Princess Margaret Hospital, Toronto, Canada; Division of Oncology, Department of Clinical Sciences, Lund University, Lund, Sweden; Departments of Cancer Epidemiology and Gastrointestinal Oncology, Moffitt Cancer Center and Research Institute, Tampa, FL; Genome Diagnostics Program, IFOM-The FIRC (Italian Foundation for Cancer Research) Institute of Molecular Oncology, Milan, Italy; Unit of Molecular Bases of Genetic Risk and Genetic Testing, Department of Research, Fondazione IRCCS Istituto Nazionale dei Tumori (INT), Milan, Italy; Adult Cancer Program, Lowy Cancer Research Centre, University of New South Wales, Sydney, Australia; The Kinghorn Cancer Centre, Garvan Institute of Medical Research, University of New South Wales, Sydney, Australia; Department of Obstetrics and Gynecology, Duke University Medical Center, Durham, NC; Department of Chronic Disease Epidemiology, Yale School of Public Health, New Haven, CT; Department of Oncology, University Hospital of Larissa, Larissa, Greece; Genomics Center, Centre Hospitalier Universitaire de Québec - Université Laval, Research Center, Québec City, Canada; Maria Sklodowska-Curie Memorial Cancer Center and Institute of Oncology, Warsaw, Poland; Department of Population and Quantitative Health Sciences, Case Western Reserve University, Cleveland, OH; Department of Health Sciences Research, Division of Biomedical Statistics and Informatics, Mayo Clinic, Rochester, MN; Division of Epidemiology, Department of Medicine, Vanderbilt Epidemiology Center, Vanderbilt-Ingram Cancer Center, Vanderbilt University School of Medicine, Nashville, TN; Department of Population Health Sciences, Huntsman Cancer Institute, Salt Lake City, UT; Division of Gynecologic Oncology, Department of Obstetrics and Gynecology, Duke University Medical Center, Durham, NC; Women’s Cancer Program at the Samuel Oschin Comprehensive Cancer Institute, Cedars-Sinai Medical Center, Los Angeles, CA; Department of Public Health Sciences, University of Virginia, Charlottesville, VA; Centre for Cancer Genetic Epidemiology, Department of Oncology, University of Cambridge, Cambridge, UK; Department of Epidemiology, Rollins School of Public Health, Emory University, Atlanta, GA

## Abstract

Familial, genome-wide association (GWAS), and sequencing studies and genetic correlation analyses have progressively unraveled the shared or pleiotropic germline genetics of breast and ovarian cancer. In this study, we aimed to leverage this shared germline genetics to improve the power of transcriptome-wide association studies (TWAS) to identify candidate breast cancer and ovarian cancer susceptibility genes. We built gene expression prediction models using the PrediXcan method in 681 breast and 295 ovarian tumors from The Cancer Genome Atlas and 211 breast and 99 ovarian normal tissue samples from the Genotype-Tissue Expression project and integrated these with GWAS meta-analysis data from the Breast Cancer Association Consortium (122,977 cases/105,974 controls) and the Ovarian Cancer Association Consortium (22,406 cases/40,941 controls). The integration was achieved through novel application of a pleiotropy-guided conditional/conjunction false discovery rate approach for the first time in the setting of a TWAS. This identified 14 new candidate breast cancer susceptibility genes spanning 11 genomic regions and 8 new candidate ovarian cancer susceptibility genes spanning 5 genomic regions at conjunction FDR < 0.05 that were > 1 Mb away from known breast and/or ovarian cancer susceptibility loci. We also identified 38 candidate breast cancer susceptibility genes and 17 candidate ovarian cancer susceptibility genes at conjunction FDR < 0.05 at known breast and/or ovarian susceptibility loci. Overlaying candidate causal risk variants identified by GWAS fine mapping onto expression prediction models for genes at known loci suggested that the association for 55% of these genes was driven by the underlying GWAS signal.

**Significance:** The 22 new genes identified by our cross-cancer analysis represent promising candidates that further elucidate the role of the transcriptome in mediating germline breast and ovarian cancer risk.

## Introduction

The last three decades have witnessed major advances in our understanding of the shared inherited genetic basis of breast and ovarian cancer. The identification of rare inherited mutations in *BRCA1* (1) and *BRCA2* (2) that confer high risks of developing both breast and ovarian cancer has directly opened up the identification of novel oncogenic mechanisms leading to the development of poly ADP ribose polymerase inhibitor therapy (3). The findings from genome-wide association studies (GWAS) have demonstrated that there is a strong genetic correlation between breast and ovarian cancer (4) and have identified several genomic regions containing common (minor allele frequency > 1%) variants that confer risk of developing both breast and ovarian cancer (5,6).

Transcriptome-wide association studies (TWAS) represent the latest study design for the identification of disease-associated susceptibility genes. TWAS involve establishing robust multi-variant models for the component of somatic (normal or tumor) gene expression that is regulated by germline genetic variation in a smaller data set where both germline genotype and somatic transcriptomic data are available. These models are then used to impute the germline genetically regulated component of gene expression into a larger GWAS data set where measured gene expression is unavailable but that offers significantly improved power to identify genes associated with disease risk where such risk may be mediated by expression. Moving from single variants (GWAS) to genes (TWAS) as the unit of association reduces the multiple testing burden. The use of gene expression provides a readily accessible read-out of the functional basis of the identified association in contrast to GWAS-identified risk variants that predominantly reside in non-coding regions of the genome (7).

PrediXcan is a method developed recently for conducting TWAS (8). TWAS methods have been applied to single cancer types before, including breast cancer (9,10) and ovarian cancer (11,12). Here we present the first application of PrediXcan, and indeed broadly of TWAS, in the pleiotropic cross-cancer setting. We used the normal and tumor breast- and ovary-specific gene expression and matched germline genotype data sets to generate tissue-specific PrediXcan models and first imputed these models into GWAS data for the corresponding cancers (i.e., from breast tissue-derived models into breast cancer GWAS and likewise for the ovarian models). We then imputed models across cancer types (i.e., from breast tissue-derived models into ovarian cancer GWAS and *vice versa*). Finally, we implemented a powerful conjunction false discovery rate approach (13,14) that has been applied previously to GWAS (15–18), but not to TWAS, to leverage the combined GWAS sample of over 145,000 breast and ovarian cancer cases. We identify new candidate breast and ovarian cancer susceptibility genes in regions not previously implicated by GWAS or TWAS analyses of these cancers.

## Methods

### Matched germline genotype – normal/tumor gene expression data sets

We used data for 211 normal breast tissue samples and 99 normal ovarian tissue samples from the Genotype-Tissue Expression (GTEx) project (version 7 release; (19)). Germline genotypes in the GTEx data had been called from whole-genome sequencing (Illumina HiSeq X) and gene expression was profiled using RNA-Sequencing (Illumina TruSeq). We also used data from 681 breast cancer (20) and 295 high-grade serous ovarian cancer [HGSOC; (21)] cases from The Cancer Genome Atlas (TCGA) network. Germline genotypes in the TCGA data had been called from genotyping arrays (Affymetrix SNP 6.0) and gene expression was profiled using RNA-Sequencing (Illumina HiSeq 2000). Imputation of TCGA germline genotypes using the 1000 Genomes version 5 reference panel was performed as described previously (22,23). The TCGA sample sizes reported here refer to only those samples that had > 95% European ancestry. Ancestry was estimated using the Local Ancestry in adMixed Populations tool (LAMP version 2.5; (24)). Downstream PrediXcan modelling (described below) used variants imputed with quality > 0.8 that had a minor allele frequency > 5% in TCGA data sets.

### Genome-wide association data sets

Summary statistics from genome-wide association meta-analyses were obtained from the Breast Cancer Association Consortium (BCAC; (22)) and the Ovarian Cancer Association Consortium (OCAC; (23)). The breast cancer susceptibility data were based on 122,977 cases and 105,974 controls, including 21,468 estrogen receptor (ER)-negative cases. The ovarian cancer susceptibility data were based on 22,406 epithelial ovarian cancer cases and 40,941 controls, including 13,037 HGSOC cases. We harmonised the signs of the effect size estimates and aligned them to the same effect allele in the breast and ovarian cancer GWAS data sets. We retained 9,530,997 variants with minor allele frequency > 1% and imputation quality > 0.4 in both data sets for S-PrediXcan analyses. All individuals in these studies were of genetically inferred European ancestry.

### PrediXcan model development and S-PrediXcan analyses

We built genetically regulated gene expression prediction models using the elastic net regularization approach implemented in PrediXcan and validated these models using tenfold cross-validation (8). Essentially, this generates a list of variants for each gene where model construction is successful and each variant in the list is assigned a weight reflecting its influence on its target gene expression. Genes with models where the nested tenfold cross-validated correlation between predicted and actual levels of expression was > 10% (predictive performance *r*^2^ > 0.01) and *P*-value of the correlation test was < 0.05 were retained. These models were adjusted for the latent determinants of gene expression variation (referred to hereafter as “PEER factors”), which were identified using the Probabilistic Estimation of Expression Residuals (PEER; version 1.3) method (25). We adjusted for 60 and 45 PEER factors for TCGA breast and ovarian cancer data, respectively. The choice of these numbers is a function of sample size and consistent with recommendations (8,25). *ESR1* expression was also included as a covariate in the construction of breast cancer models to account for estrogen receptor status and its influence on the expression of individual genes. For the GTEx version 7 data sets, we downloaded pre-computed PrediXcan models from predictdb.org. Our pipeline for processing the TCGA data sets, including the application of PEER factors, was designed to be consistent with the pipeline used to generate the pre-computed GTEx PrediXcan models. S-PrediXcan refers to the application of the PrediXcan gene expression models, specifically the variant weights from elastic net combined into multi-variant gene-level instruments, to summary statistics GWAS data sets and has been described in detail before (8).

### Conditional and conjunction false discovery rate analyses

We obtained *P*-values for association of predicted expression of each gene with breast cancer risk and with ovarian cancer risk. We then computed the false discovery rate (FDR) for gene-breast cancer risk association conditional on gene-ovarian cancer risk association (as conditional FDR_Breast Cancer|Ovarian Cancer_). This is the probability that a gene is not associated with breast cancer risk given the P-values for association with both breast cancer risk and ovarian cancer risk. The analogous conditional FDR for gene-ovarian cancer risk association was also calculated (FDR_Ovarian Cancer|Breast Cancer_). Finally, the conjunctional FDR estimate, which is conservatively defined as the maximum of the two conditional FDR values, was computed. This process minimizes the effect of a single phenotype (in this case, breast or ovarian cancer) driving the shared association signal. It allows the power of pleiotropic associations to be tapped for genetic discovery unlike a traditional FDR approach that is informed solely by the distribution of *P*-values for a single phenotype. We used the R implementation of the conditional FDR method available from github.com/KehaoWu/GWAScFDR. The conditional and conjunctional FDR method has been described extensively elsewhere (13–18), but not applied before to the TWAS setting. The overall study design is summarized in Figure 1.

**Figure 1:**
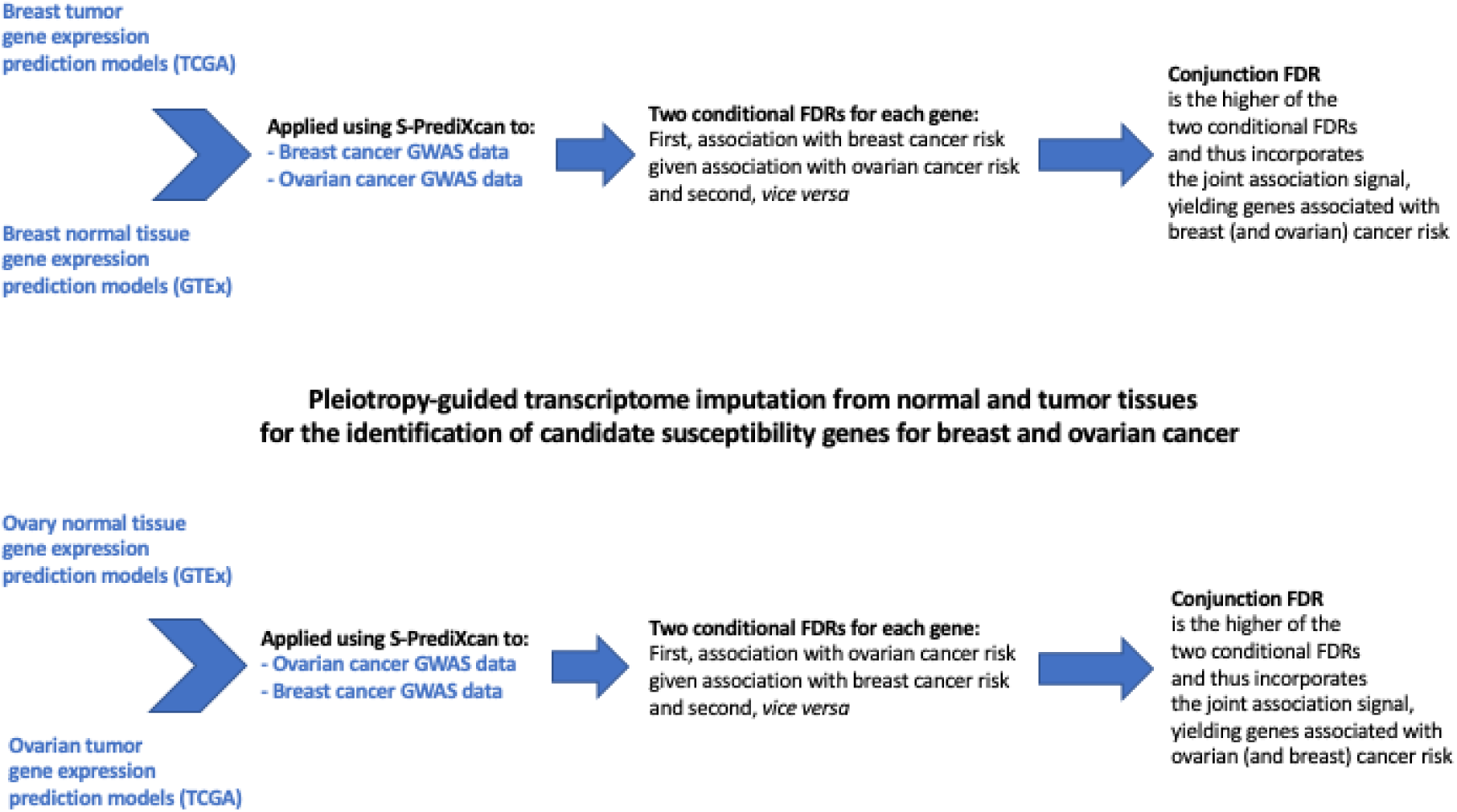
Flowchart providing an overview of the data sets used and the various steps in the analysis. GTEx: Genotype-Tissue Expression project; TCGA: The Cancer Genome Atlas; GWAS: genome-wide association study; FDR: false discovery rate.

### Fine-mapped candidate causal risk variant data sets

We examined the overlap between variants in the breast gene expression prediction models and a published list of fine-mapped candidate causal risk variants for breast cancer (26). This was done to follow-up genes that we identified in genomic regions that are known to be associated with breast cancer risk under the intuition that gene-level association signals identified by S-PrediXcan that demonstrate such overlap with fine-mapped variants are likely being driven by the GWAS association signal in the same region.

Fine-mapped candidate causal risk variants lists for breast cancer were obtained from Fachal et al (26). Briefly, Fachal et al. fine-mapped 150 known breast cancer susceptibility regions using dense genotype data on women participating in the BCAC and in the Consortium of Investigators of Modifiers of BRCA1/2 (CIMBA). Stepwise multinomial logistic regression was used to identify independent association signals in each region. Credible causal variants within each signal were defined as being within a 100-fold likelihood of the top conditional variant to delineate the variants driving the GWAS associations in each region.

## Results

### Development of tissue/tumor-specific gene expression prediction models

We built genetically regulated gene expression predictor models using matched germline genotype and tumor gene expression data from TCGA by applying elastic net regularization as implemented in the PrediXcan software. Genes with models where the nested tenfold cross-validated correlation between predicted and actual levels of expression was > 10% (predictive performance *r*^2^ > 0.01) and *P*-value of the correlation test was < 0.05 were retained in line with best practice quality control recommendations by the developers of PrediXcan (8). We constructed and evaluated predictor models that met these criteria for 4,457 genes based on 681 TCGA breast tumor samples and for 2,705 genes based on 295 TCGA ovarian tumor samples. We obtained pre-computed genetically regulated gene expression predictor models that met the same criteria (predictive performance *r*^2^ > 0.01; correlation test *P* < 0.05) in matched germline genotype and normal tissue gene expression data from the GTEx Project. Specifically, the pre-computed data included 5,274 genes modelled based on 211 GTEx breast tissue samples and 3,034 genes modelled based on 99 GTEx ovarian tissue samples.

### Imputation of gene expression into GWAS and pleiotropy-guided FDR control

We used the GTEx normal breast tissue-derived prediction models to impute genetically regulated gene expression in a genome-wide association meta-analysis involving 122,977 breast cancer cases and 105,974 controls using S-PrediXcan. We tested for association between imputed gene expression and breast cancer risk. We also used the same GTEx breast tissue-based models to impute gene expression in a genome-wide association meta-analysis of 22,406 ovarian cancer cases and 40,941 controls and test for association between imputed expression and ovarian cancer risk. For these two steps, we applied the conditional FDR method to the S-PrediXcan gene-level association *P*-values to correct for testing 5,274 genes in each analysis. This yielded two conditional FDR values: one for association with breast cancer risk given association with ovarian cancer risk and the other for association with ovarian cancer risk given association with breast cancer risk. Finally, we took the larger of the two values for each gene as a conservative estimate of its conjunction FDR to identify candidate breast cancer susceptibility genes at conjunction FDR < 0.05. We refer to these genes as candidate breast cancer susceptibility genes because they were identified on the basis of gene expression predictor models derived from breast tissue. However, the conditional-conjunction FDR analysis effectively borrowed information from pleiotropic associations with inherited susceptibility to a second cancer type (in this case ovarian cancer) in addition to the primary cancer type (breast cancer) and these genes may be considered as risk genes for the second cancer as well. These steps were repeated for three other ordered combinations of data sets: TCGA breast tumor tissue-breast cancer GWAS-ovarian cancer GWAS to identify candidate breast cancer susceptibility genes; GTEx normal ovarian tissue-ovarian cancer GWAS-breast cancer GWAS and TCGA ovarian tumor tissue-ovarian cancer GWAS-breast cancer GWAS to identify candidate ovarian cancer susceptibility genes. We also replaced the overall breast cancer GWAS and all invasive ovarian cancer GWAS used in the four data set combinations described above with ER-negative breast cancer GWAS (21,468 cases/105,974 controls) and HGSOC GWAS (13,037 cases/22,406 controls), respectively. This helped identify additional candidate breast and ovarian cancer susceptibility genes driven by subtype-specific associations at conjunction FDR < 0.05.

For each gene, coverage was defined as the percentage of the number of variants included in its expression prediction model that were also captured in the genome-wide association meta-analysis. The coverage was >= 80% for at least 93% of the genes in each of the four matched germline genotype and normal or tumor gene expression data sets used to build the predictor models indicating that for most genes, most of the corresponding model variants available were used. In each ordered analytic combination of data sets (e.g., GTEx normal breast tissue-breast cancer GWAS-ovarian cancer GWAS) we observed that, in general, for progressively smaller S-PrediXcan *P*-values of the second cancer type, the true discovery rate for association with the primary cancer type approached 100% at progressively larger S-PrediXcan *P*-values for the primary cancer type (Figure 2 and Supplementary Figure 1). This was consistent with substantial shared gene-level associations for breast and ovarian cancer risk and these shared signals being tapped by the conditional-conjunction FDR method to power candidate susceptibility gene discovery.

**Figure 2:**
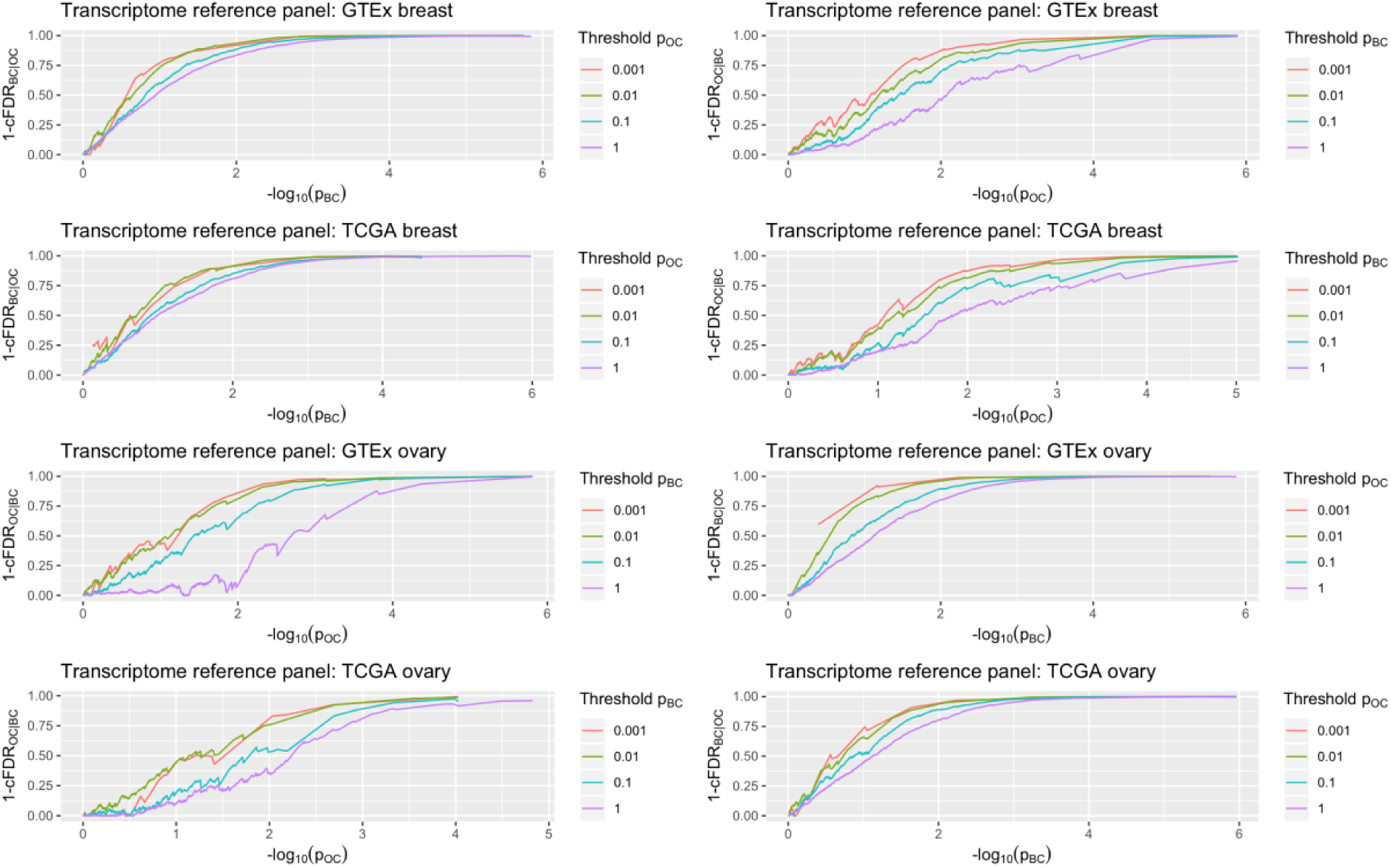
True discovery rate against the negative logarithm (base 10) of the *P*-value for each cancer for subsets of genes based on strength of association with the other cancer. The Y-axis of each plot is the true discovery rate which is defined as 1 – conditional false discovery rate (cFDR). For a given ordered analytic combination of data sets (e.g., GTEx normal breast tissue as transcriptome reference panel-breast cancer GWAS-ovarian cancer GWAS, plotted in the upper left hand corner) we observed that, in general, for progressively smaller S-PrediXcan *P*-values of the second cancer type (indicated by the key “Threshold p” next to each plot), the true discovery rate (Y-axis) for association with the primary cancer type approached 100% at progressively larger S-PrediXcan *P*-values for the primary cancer type (X-axis; negative logarithm (base 10) of the *P*-values). BC: overall breast cancer risk; OC: all invasive ovarian cancer risk. Only *P*-values > 10^−6^ are plotted on the X-axis.

### Identification of new candidate breast cancer and ovarian cancer susceptibility genes

We identified 14 new candidate breast cancer susceptibility genes at the conjunction FDR < 0.05 threshold (Table 1 and Supplementary Table 1). The 14 genes were distributed between 11 genomic regions > 1 Mb apart from each other (Table 1). These genes have not been reported as susceptibility genes in any prior TWAS of breast cancer risk and are > 1 Mb away from published genome-wide significant lead variants for breast cancer susceptibility (27). For ovarian cancer, we identified 8 novel candidate susceptibility genes at conjunction FDR < 0.05 (Table 2 and Supplementary Table 2). The 8 genes were located across 5 genomic regions > 1 Mb apart from each other (Table 2). These genes have not been reported as candidate risk genes in any previously reported TWAS of ovarian cancer risk and are > 1 Mb away from published genome-wide significant lead variants for ovarian cancer susceptibility (23).

**Table 1:** New candidate breast cancer susceptibility genes identified by pleiotropy-guided S-PrediXcan analysis.

| Gene | Genomic Region | P-value<br>BC | P-value<br>OC | Conditional FDR<br>BC\|OC | Conditional FDR<br>OC\|BC | Conjunction<br>FDR |
| --- | --- | --- | --- | --- | --- | --- |
| **Transcriptome reference panel: GTEx breast (normal) \| primary GWAS: overall BC risk (second GWAS: all invasive OC risk)** |  |  |  |  |  |  |
| ZSCAN29 | 15q15.3 | 1.8E-04 | 9.1E-04 | 4.1E-04 | 6.0E-03 | 6.0E-03 |
| STRCP1 | 15q15.3 | 1.6E-03 | 1.8E-03 | 4.1E-03 | 1.9E-02 | 1.9E-02 |
| AC011330.5 | 15q15.3 | 5.4E-04 | 2.5E-03 | 1.8E-03 | 2.2E-02 | 2.2E-02 |
| STRC | 15q15.3 | 1.4E-04 | 3.8E-03 | 6.1E-04 | 2.2E-02 | 2.2E-02 |
| ZNF276 | 16q24.3 | 3.7E-06 | 4.7E-03 | 2.4E-05 | 2.2E-02 | 2.2E-02 |
| RGS19 | 20q13.33 | 1.1E-03 | 5.4E-03 | 4.3E-03 | 4.3E-02 | 4.3E-02 |
| RNFT1 | 17q23.1 | 2.4E-04 | 8.7E-03 | 1.3E-03 | 4.7E-02 | 4.7E-02 |
| C15orf65 | 15q21.3 | 2.2E-03 | 5.9E-03 | 8.2E-03 | 4.7E-02 | 4.7E-02 |
| **Transcriptome reference panel: TCGA breast (tumor) \| primary GWAS: overall BC risk (second GWAS: all invasive OC risk)** |  |  |  |  |  |  |
| GMNC | 3q28 | 2.6E-03 | 1.2E-03 | 6.1E-03 | 2.0E-02 | 2.0E-02 |
| ESRP2 | 16q22.1 | 1.9E-02 | 9.6E-04 | 4.3E-02 | 3.3E-02 | 4.3E-02 |
| BHLHA15 | 7q21.3 | 8.5E-05 | 7.0E-03 | 5.5E-04 | 4.9E-02 | 4.9E-02 |
| SCGB1D2 | 11q12.3 | 3.5E-04 | 5.5E-03 | 2.0E-03 | 4.9E-02 | 4.9E-02 |
| **Transcriptome reference panel: TCGA breast (tumor) \| primary GWAS: ER-negative BC risk (second GWAS: HGSOC risk)** |  |  |  |  |  |  |
| ETAA1 | 2p14 | 3.0E-03 | 1.5E-03 | 2.0E-02 | 2.0E-02 | 2.0E-02 |
| ATP8B4 | 15q21.2 | 1.6E-03 | 2.2E-03 | 1.5E-02 | 2.4E-02 | 2.4E-02 |

Abbreviations: BC, breast cancer; OC, ovarian cancer; FDR, false discovery rate; ER, estrogen receptor; HGSOC, high-grade serous ovarian cancer.

**Table 2:**
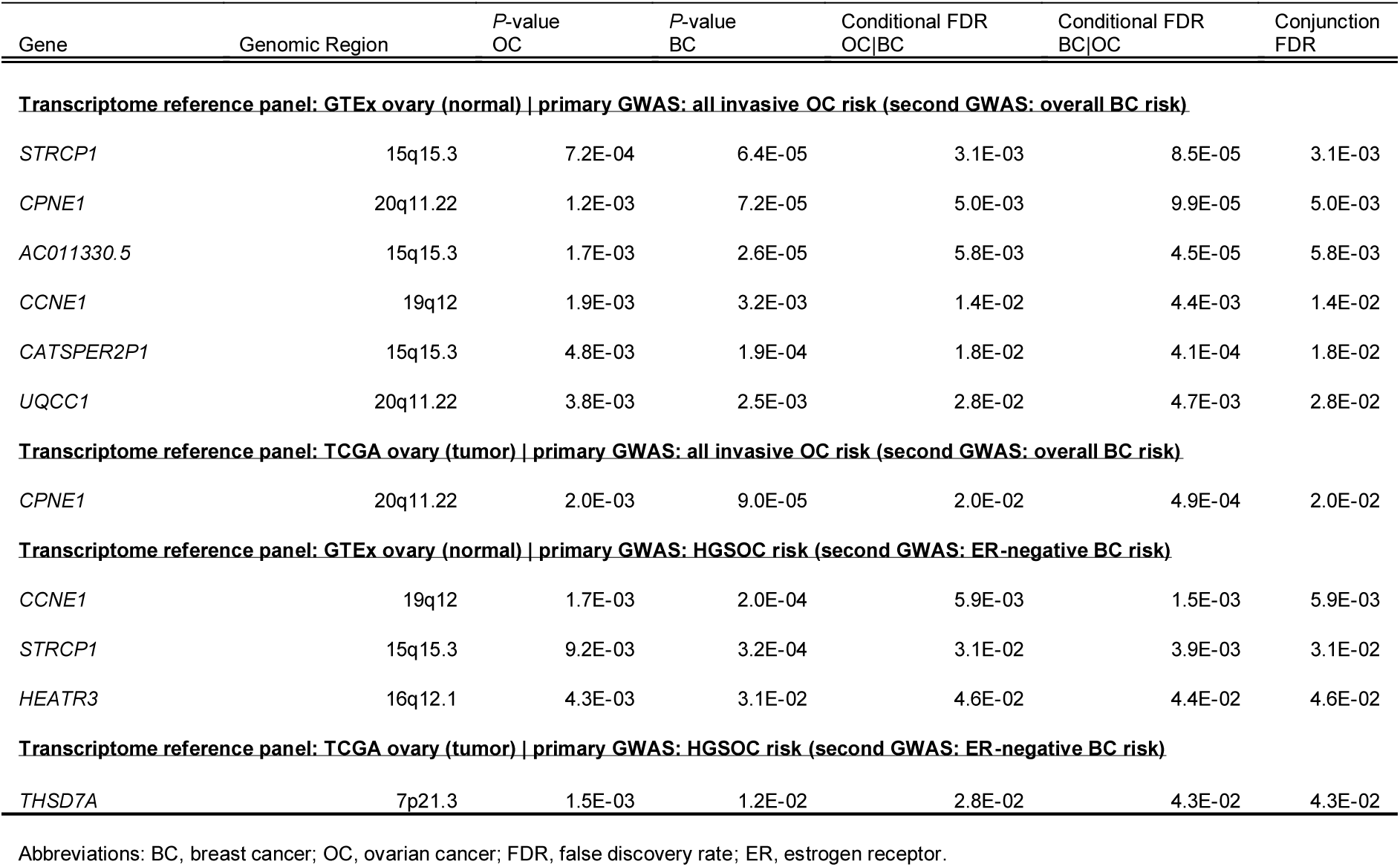
New candidate ovarian cancer susceptibility genes identified by pleiotropy-guided S-PrediXcan analysis.

### Candidate breast cancer and ovarian cancer susceptibility genes at known GWAS loci

We identified 38 candidate breast cancer susceptibility genes that were located within 1 Mb of a published lead variant associated at genome-wide significance with breast cancer risk (Supplementary Table 3; (27)). Four of the 38 genes have also been reported in previously published TWAS (Supplementary Table 3; (9,10)). The 38 genes were spread across 12 genomic regions > 1 Mb apart from each other. Overlaying fine-mapped candidate causal breast cancer risk variants on breast gene expression predictor model variants showed that for 21/38 (55%) genes the prediction model variants included at least one fine-mapped candidate causal variant (Supplementary Tables 3 and 4). This suggested that for these genes the GWAS association signal was driving the S-PrediXcan signal. We also identified three additional genes that were > 1 Mb away from known GWAS loci that have previously been reported as TWAS loci for breast cancer risk (Supplementary Table 3; (9,10)).

For ovarian cancer, we identified 17 candidate susceptibility genes that were located within 1 Mb of a published lead variant associated at genome-wide significance with ovarian cancer risk (Supplementary Table 5; (23)). Six of these genes have also been reported in a previously published TWAS for ovarian cancer (Supplementary Table 5; (11,12)). The 17 genes span 5 different genomic regions > 1 Mb apart.

## Discussion

In this study, we used the conditional and conjunctional FDR as a novel tool to systematically improve the power of breast cancer and ovarian cancer candidate susceptibility gene discovery in a PrediXcan-based TWAS. While gene expression prediction models based on multiple tissue types has been the more common approach to improving TWAS power (11,28), the conditional/conjunction FDR approach gains power through the incorporation of multiple related GWAS data sets into a TWAS analysis. We investigated the shared inherited genetic basis of these two cancer types by integrating normal and tumor tissue-specific transcriptomic data sets with large-scale genome-wide association meta-analysis findings for susceptibility to breast cancer and ovarian cancer. We identified 11 new genomic regions associated with breast cancer risk and five new regions linked to ovarian cancer risk.

We identified 14 novel candidate breast cancer susceptibility genes (Table 1). Many of these genes have a strong biological rationale for involvement in breast carcinogenesis and are in or near genomic regions associated with other cancer types or potential cancer risk factors. For example, the *ZNF276* intronic variant rs12925026 is associated at genome-wide significance with non-melanoma skin cancer (29). *ZNF276* overlaps *FANCA* in a tail-to-tail manner (30). The genetically regulated predictor model for *ZNF276* expression was fit using gene expression measured in GTEx breast tissues but neither this data set nor any of the other data sets could capture a predictor model for *FANCA* expression. *FANCA* encodes one of eight subunits that together form the core Fanconi Anemia (FA) complex that repairs blockages in DNA replication due to cross-linking (31). Several members of the FA family of proteins have been implicated in breast and ovarian cancer predisposition including, *BRCA1* (*FANCS*), *BRCA2* (*FANCD1*), *BRIP1* (*FANCJ*), *PALB2* (*FANCN*), *RAD51C* (*FANCO*) and it is possible that *FANCA* may represent another or possibly the true target breast cancer susceptibility gene in this region given this biological function and its overlap with *ZNF276* (31,32). *ZNF276* in its own right has also been implicated as a candidate tumor suppressor gene in breast cancer (30).

Other candidate breast cancer susceptibility genes we identified include *ESRP2*, which encodes an epithelial cell-specific regulator of splicing of the breast cancer susceptibility gene *FGFR2* (33,34) and *SCGB1D2*, which encodes lipophilin B that is known to be expressed in both breast and ovarian tumors (35). Lipophilin B is tightly co-expressed with and forms a covalent complex with Mammaglobin A encoded by *SCGB2A2*, the gene next to *SCGB1D2* (35). Mammaglobin A may be used to detect disseminated or circulating tumor cells and is under investigation as a potential immunotherapeutic target in breast cancer (36). However, we were unable to develop gene expression prediction models for *SCGB2A2* in breast normal or tumor tissues. *BHLHA15* encodes an estrogen-regulated transcription factor that is required to maintain mammary gland differentiation in mice (37). *ETAA1* harbors lead variants associated at genome-wide significance with pancreatic cancer (38), and the hormone-related traits of age at menopause (39) and male-pattern baldness (40). It encodes an activator of ATR kinase that accumulates at DNA damage sites and promotes replication fork progression and integrity (41). While our pleiotropy-guided transcriptome imputation study was ongoing, a genome-wide association meta-analysis for breast cancer risk that was performed in parallel identified lead variants rs79518236 (184 kb from *BHLHA15*) and rs9712235 (244 kb from *ETAA1*) at genome-wide significance only on addition of 10,407 breast cancer cases and 7,815 controls to the Michailidou et al. data set used here ((42), unpublished pre-print). There were no known GWAS associations for breast cancer risk in these regions until the larger GWAS meta-analysis and our concomitant identification of the same regions using gene expression imputation into a smaller GWAS underscores the power of leveraging expression data to bolster genetic discovery.

We identified 11 new candidate ovarian cancer susceptibility genes (Table 2). As with breast cancer, there is strong support for a role of several genes in ovarian cancer pathogenesis and many of these genes are in regions of the genome that harbor pleiotropic associations with other cancer types. Variants immediately upstream of *CCNE1* are associated at genome-wide significance with bladder cancer risk (43). *CCNE1* amplification is believed to be an early event in the development of ovarian cancer (44) and is a frequent somatic event in HGSOCs that do not carry homologous recombination DNA repair pathway defects (45). *CCNE1* amplification is also associated with poor prognosis in triple negative breast tumors (46) and it is worth noting that we observed the stronger conjunction FDR association signal for *CCNE1* in the pleiotropy-informed analysis that was based on the HGSOC and ER-negative breast cancer susceptibility GWAS data sets (Table 2). This study is the first to suggest a role for *CCNE1* in conferring ovarian cancer risk. Intronic variants in *HEATR3* are associated at genome-wide significance with glioma in European ancestry individuals (47) and with squamous cell esophageal carcinoma in East Asian ancestry individuals (48). *HEATR3* was also identified by a TWAS of glioma susceptibility (49). Intronic variants in *THSD7A* are associated with epithelial ovarian cancer risk in East Asians (50), albeit not at genome-wide significance (lead variant rs10260419 *P* = 1 × 10^−7^). Gene expression prediction models derived from breast and ovarian tissues both implicated the 15q15.3 region as a new breast and ovarian cancer susceptibility region on imputation with these models into the breast and ovarian cancer GWAS data. Our analysis suggested several genes in this region (Tables 1 and 2), with the pseudogene *STRCP1* as the only common gene across breast and ovarian tissues. *STRCP1* overlaps the protein coding gene *STRC*, also identified in the breast tissue-based analysis (Table 1), and variants in *STRC* have previously been associated with lung cancer risk (lung cancer lead variant rs35028925 *P* = 2 × 10^−6^) (51).

In this analysis, we chose to label the identified genes as candidate breast cancer susceptibility genes if they were identified on integrating the GTEx or TCGA breast expression prediction models with the breast cancer GWAS and incorporating pleiotropic information from the ovarian cancer GWAS and *vice versa* for candidate ovarian cancer susceptibility genes. However, application of the conjunction FDR over and above the conditional FDR in principle identified genes associated with both cancer types by tapping into GWAS data from both cancers. Therefore, in a sense all these genes may well be regarded as candidate breast *and* ovarian cancer susceptibility genes.

We identified 38 candidate breast cancer susceptibility genes and 17 candidate ovarian cancer susceptibility genes in regions previously implicated by GWAS for breast cancer and ovarian cancer, respectively (Supplementary Tables 3 and 5). The identification of a large number of genes in these regions is unsurprising given that GWAS associations are the key determinant of the S-PrediXcan signal. However, we were able to take full advantage of extensive fine-scale mapping data generated by the Breast Cancer Association Consortium to separately pinpoint those genes where a fine-mapped candidate causal GWAS risk variant for breast cancer was incorporated in the breast PrediXcan model suggesting it drives the gene-based association. Overall, we found this to be the case for 55% of the candidate susceptibility genes identified by PrediXcan in the breast cancer susceptibility regions identified by GWAS. Comprehensive functional follow up of the 19p13.11 breast and ovarian cancer GWAS region suggests that *ABHD8* and *ANKLE1* are the most likely targets in this region (5). While there was no overlap between S-PrediXcan model variants for *ABHD8* and *ANKLE1* and fine-mapped risk variants in this region, S-PrediXcan did detect both genes as candidate causal susceptibility genes, with *ANKLE1* being the only gene that made the cut in both breast and ovarian tissues, suggesting that S-PrediXcan applied to pleiotropic gene dense regions such as 19p13.11 does help short-list the key targets even in the absence of overlap with fine-mapped variants. A total of 21/38 breast and 13/17 ovarian cancer candidate susceptibility genes in the published GWAS regions were clustered at 17q21.31, reflecting the unique long-distance linkage disequilibrium structure of this region (52). This phenomenon has also led to clustering of associations at 17q21.31 in previous TWAS of breast or ovarian cancer risk (9,11).

In conclusion, the powerful combination of pleiotropic breast and ovarian cancer GWAS data sets with transcriptome imputation from normal and tumor breast and ovarian tissues identified a total of 16 novel genomic loci (22 new genes) associated with breast and ovarian cancer risks. Fine-mapping in larger GWAS data sets and deeper laboratory-based functional follow-up studies of these new loci and candidate genes have the potential to provide fresh insights into the common biological underpinnings of breast and ovarian cancer.

## Data Access Statement

Genome-wide summary genetic association statistics from BCAC are available at: http://bcac.ccge.medschl.cam.ac.uk/bcacdata/oncoarray/gwas-icogs-and-oncoarray-summary-results/

Genome-wide summary genetic association statistics from OCAC are available at: https://www.ebi.ac.uk/gwas/downloads/summary-statistics

PrediXcan prediction models trained on the GTEx version 7 data (breast and ovarian tissues) are available here: http://hakyimlab.org/post/2017/v7-v6p-analysis/

PrediXcan prediction models trained on the TCGA data (breast and ovarian tumors) will be made publicly available upon publication.

## Supporting information

All Supplementary Tables

All Supplementary Figures

## Acknowledgments

The analyses presented in this manuscript were funded by grant number R01CA211574 from the United States National Institutes of Health/National Cancer Institute.

## For GTex

The GTEx Project was supported by the Common Fund of the Office of the Director of the National Institutes of Health, and by NCI, NHGRI, NHLBI, NIDA, NIMH, and NINDS. The data used for the analyses described in this manuscript can be obtained from dbGaP via accession number phs000424.

## For TCGA

The results published here are in part based upon data generated by the TCGA Research Network: https://www.cancer.gov/tcga.

## For BCAC

The breast cancer genome-wide association analyses were supported by the Government of Canada through Genome Canada and the Canadian Institutes of Health Research, the ‘Ministère de l’Économie, de la Science et de l’Innovation du Québec’ through Genome Québec and grant PSR-SIIRI-701, The National Institutes of Health (U19 CA148065, X01HG007492), Cancer Research UK (C1287/A10118, C1287/A16563, C1287/A10710) and The European Union (HEALTH-F2-2009-223175 and H2020 633784 and 634935). All studies and funders are listed in Michailidou et al (2017).

## For OCAC

The ovarian cancer genome-wide association meta-analyses were supported by the US National Institutes of Health (CA1×01HG007491-01 (C.I. Amos), U19-CA148112 (T.A. Sellers), R01-CA149429 (C.M. Phelan) and R01-CA058598 (M.T. Goodman); Canadian Institutes of Health Research (MOP-86727 (L.E. Kelemen)) and the Ovarian Cancer Research Fund (A. Berchuck). The COGS project was funded through a European Commission’s Seventh Framework Programme grant (agreement number 223175 - HEALTH-F2–2009-223175). All studies and funders are listed in Phelan et al (2017).

