## Supplementary material for "Pleiotropy-guided transcriptome imputation from normal and tumor tissues identifies new candidate susceptibility genes for breast and ovarian cancer": All Supplementary Figures

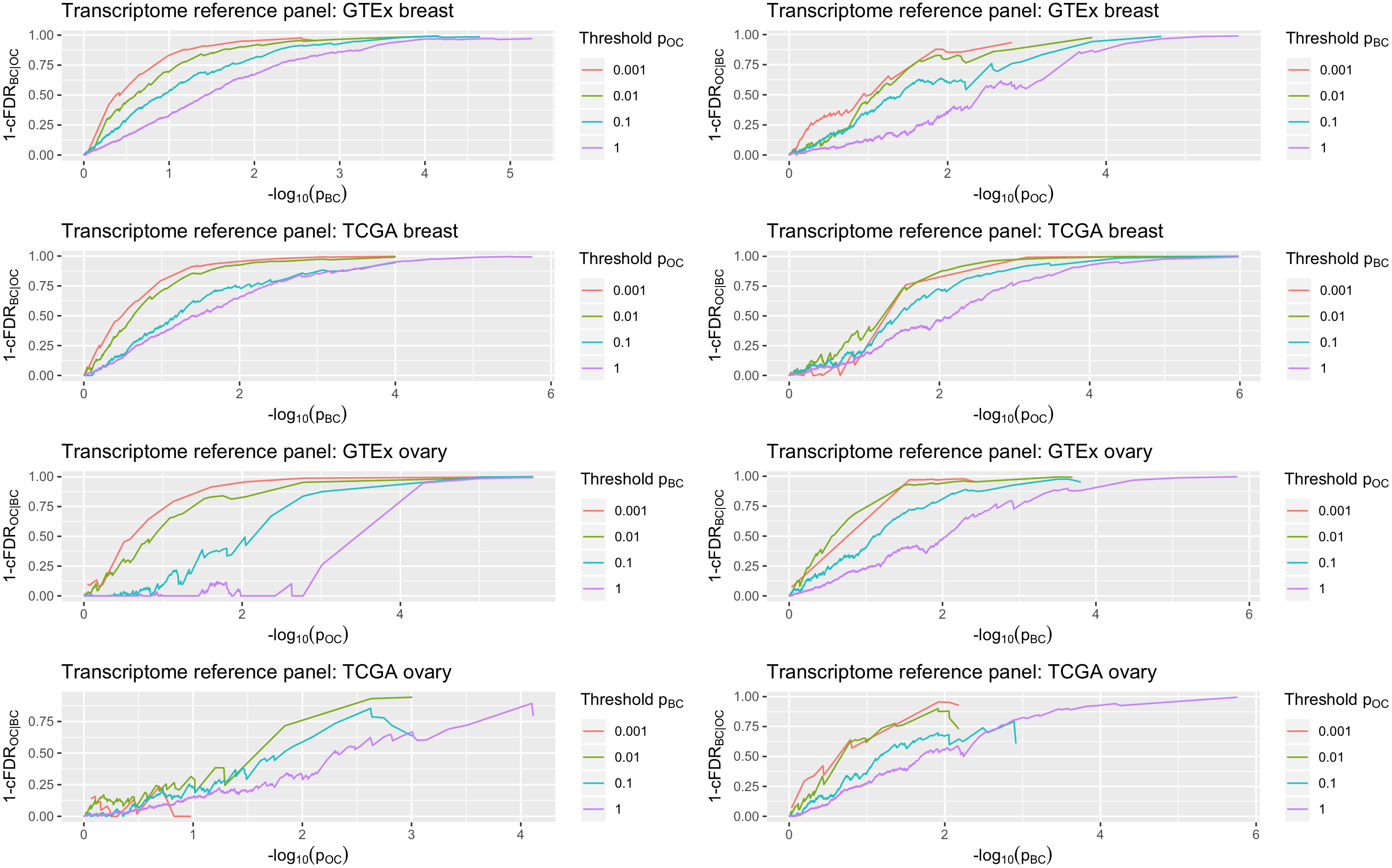

**Supplementary Figure 1:** True discovery rate against the negative logarithm (base 10) of the *P*-value for each cancer for subsets of genes based on strength of association with the other cancer for the subtype-specific analyses. The Y-axis of each plot is the true discovery rate which is defined as 1 – conditional false discovery rate (cFDR). For a given ordered analytic combination of data sets (e.g., GTEx normal breast tissue as transcriptome reference panel-estrogen receptor (ER)-negative breast cancer GWAS-high-grade serous ovarian cancer (HGSOC) GWAS, plotted in the upper left hand corner) we observed that, in general, for progressively smaller S-PrediXcan *P*-values of the second cancer type (indicated by the key “Threshold p” next to each plot), the true discovery rate (Y-axis) for association with the primary cancer type approached 100% at progressively larger S-PrediXcan *P*-values for the primary cancer type (X-axis; negative logarithm (base 10) of the *P*-values). BC: ER-negative breast cancer risk; OC: HGSOC risk. Only *P*-values > 10^-6^ are plotted on the X-axis.
